## Supplementary Materials for "nnSVG for the scalable identification of spatially variable genes using nearest-neighbor Gaussian processes"

### Contents

1. **Supplementary Figures S1-S24**

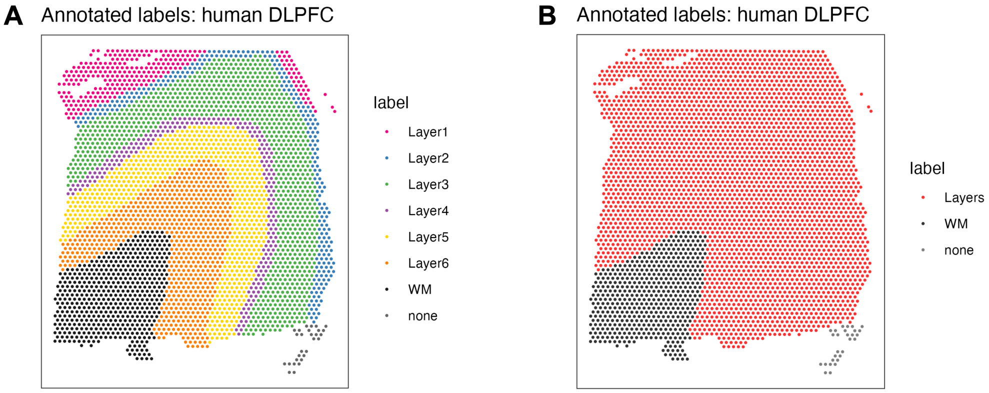


**Supplementary Figure S1: Manually annotated layer labels for Visium human DLPFC dataset.** **(A)** Manually annotated cortical layer labels for Visium human DLPFC dataset, sourced from [8], which we use as an approximate ground truth for method evaluation. The DLPFC contains six cortical layers and white matter. **(B)** Same as A, comparing cortical layers and white matter only. The most biologically significant differences in expression in this dataset are between the cortical layers and white matter [8]. WM = white matter.

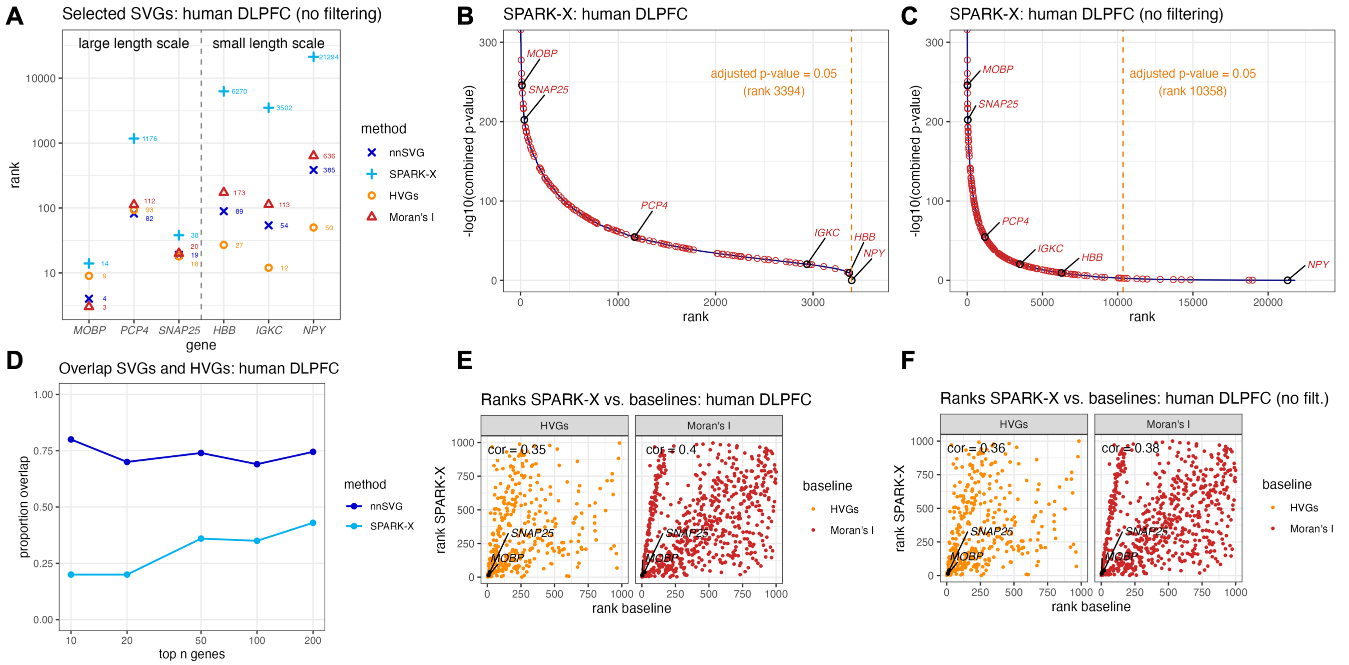


**Supplementary Figure S2: Additional results for Visium human DLPFC dataset.** **(A)** Rank order of the 6 SVGs from Figure 1 within the lists of top SVGs for nnSVG, SPARK-X, HVGs, and Moran's I, without filtering low-expressed genes for nnSVG or SPARK-X. Dashed vertical line divides the genes into the 3 cortical layer-associated SVGs with large length scales (left) and the 3 blood- and immune-associated SVGs with small length scales (right). **(B)** Combined $p$-value from SPARK-X ($y$-axis) compared to the rank per gene ($x$-axis), with the 6 SVGs from (A) labeled, and 134 additional known layer-specific marker genes (from manually guided analyses by [8]) highlighted (red circles). Orange dashed vertical line indicates rank cutoff for statistically significant SVGs at a multiple-testing-adjusted p-value of 0.05 using statistical test defined in [27]. **(C)** Same as (B), without filtering low-expressed genes. **(D)** Overlap between sets of top $n$ SVGs and HVGs ($n = 10, 20, 50, 100, 200$) for nnSVG and SPARK-X, for main results including filtering of low-expressed genes. **(E)** Ranks of top 1000 SVGs from SPARK-X ($y$-axis) compared to ranks from baseline methods ($x$-axis) using HVGs (nonspatial baseline method, left) and Moran's I (spatially-aware baseline method, right), with SVGs from (A) highlighted (black circles), and Spearman correlation (text labels). **(F)** Same as (E) for SPARK-X, without filtering low-expressed genes. For the results without filtering low-expressed genes, we used parameters `order = Sum_coords` and `n.neighbors = 15` for nnSVG for improved stability for low-expressed genes.

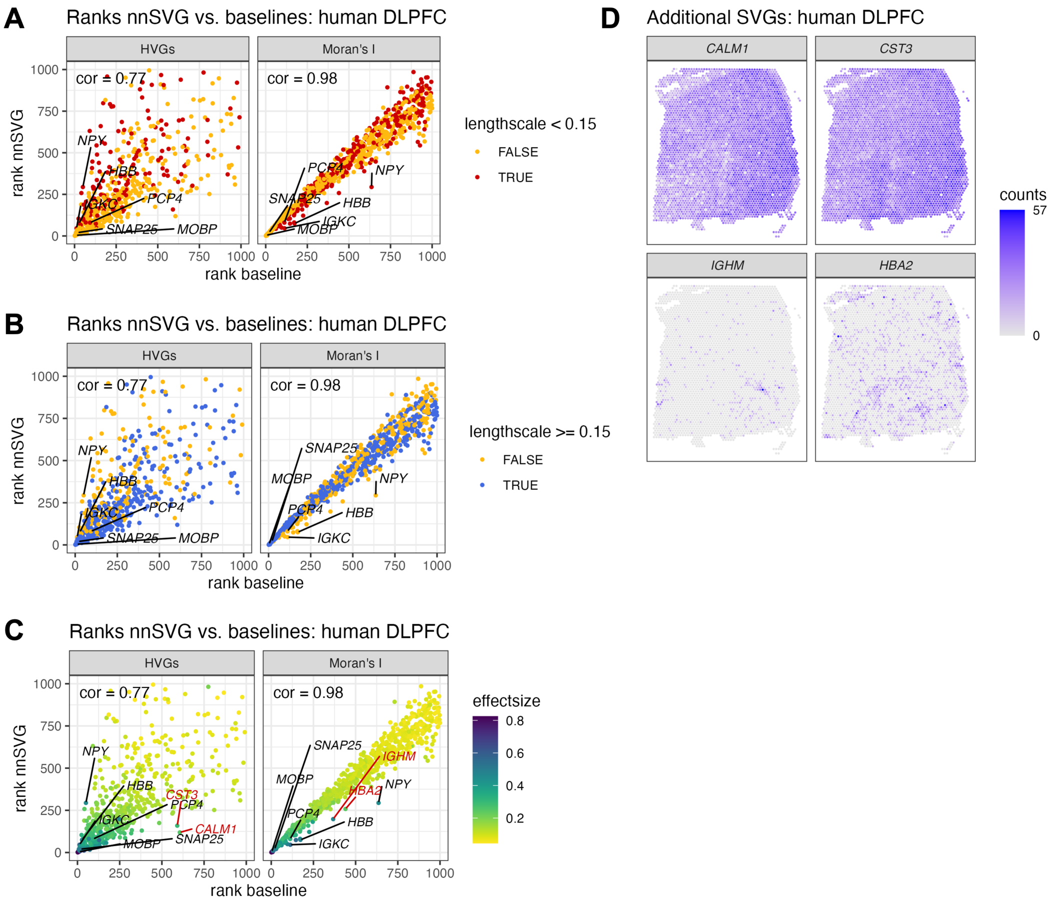


**Supplementary Figure S3: Additional results for Visium human DLPFC dataset.** **(A–B)** Ranks of top 1000 SVGs from nnSVG compared to baseline methods (HVGs and Moran's I, left and right panels respectively), highlighting genes with estimated length scale parameters $< 0.15$ (red, A) and $>= 0.15$ (blue, B). **(C)** Ranks of top 1000 SVGs from nnSVG compared to baseline methods (HVGs and Moran's I, left and right panels respectively), with color scale indicating effect size estimates. The 6 known SVGs from the main results are labeled, along with 2 additional SVGs in the lower-right off-diagonal quadrant (red labels) in each panel. **(D)** Spatial expression plots for the 4 additional SVGs highlighted in C.

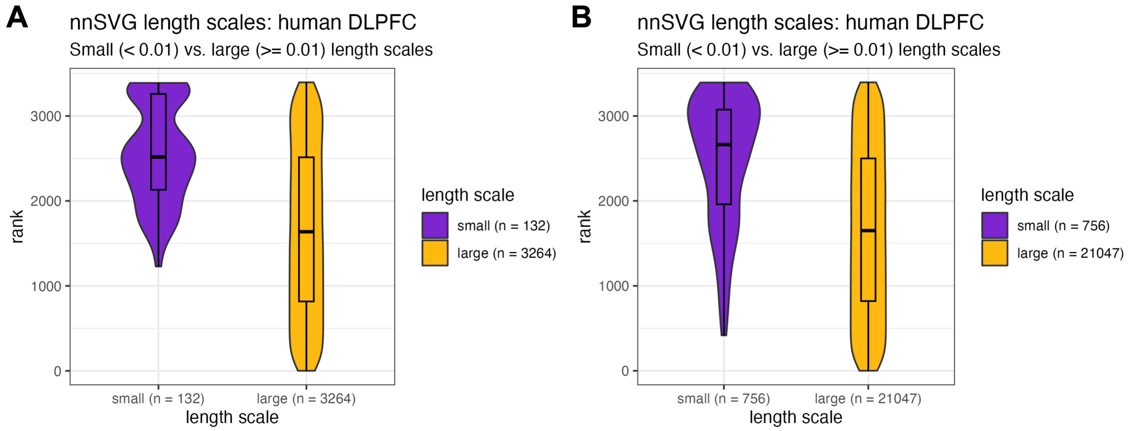


**Supplementary Figure S4: Additional results for Visium human DLPFC dataset.** **(A)** Rank distributions for genes with extremely small estimated length scales (< 0.01) in the results with filtering to remove low-expressed genes (consistent with the main results). **(B)** Rank distributions for genes with extremely small estimated length scales (< 0.01) without filtering to remove low-expressed genes (showing top 3,396 genes only, i.e. the same number of genes as in A, for comparability). Legends indicate the number of genes $n$ within each group. Boxplots show medians, first and third quartiles, and whiskers extending to the furthest values no more than 1.5 times the interquartile range from each quartile.

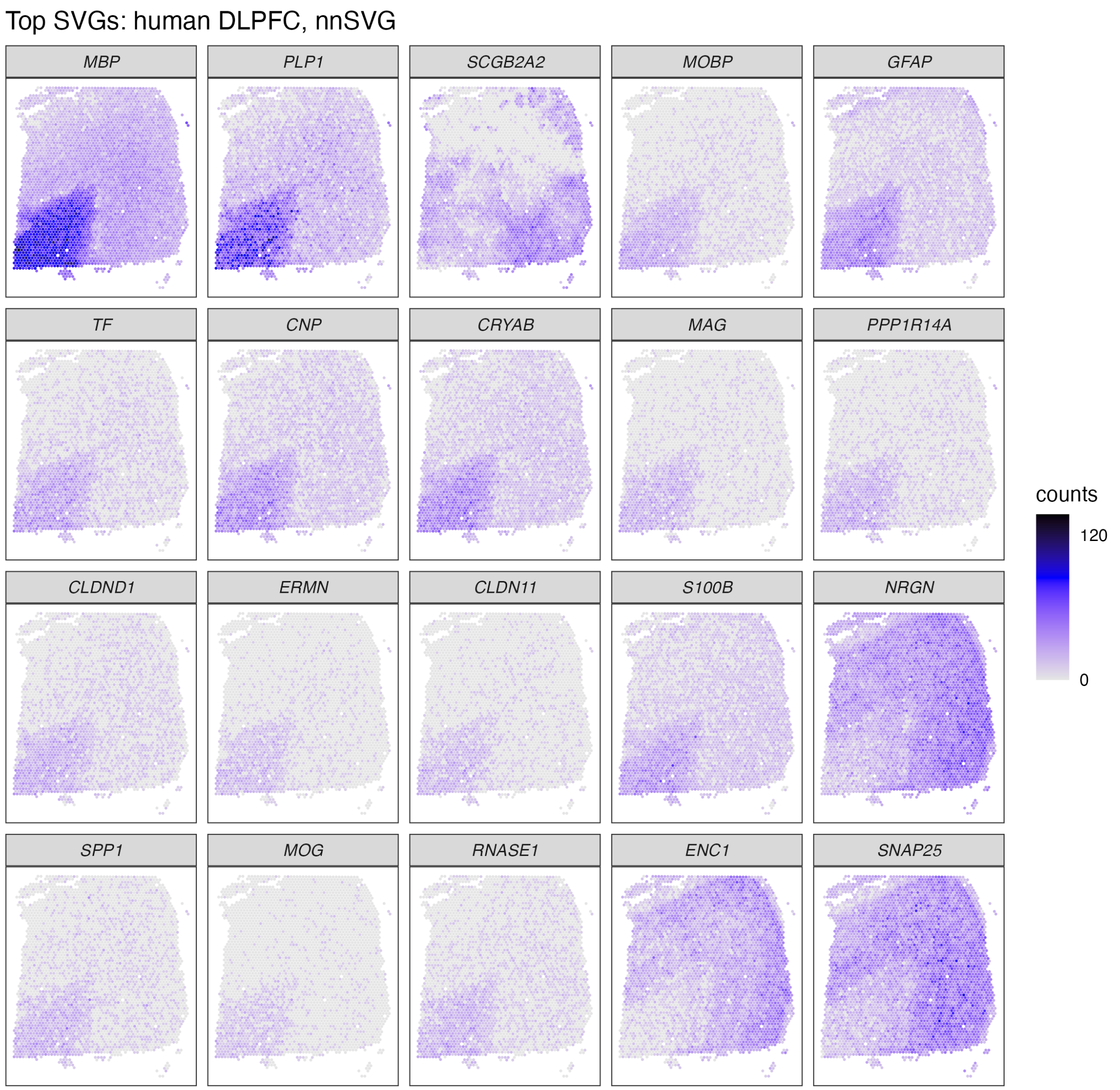


**Supplementary Figure S5: Spatial expression of top SVGs, Visium human DLPFC, nnSVG.** Spatial expression plots of top 20 SVGs from nnSVG for Visium human DLPFC dataset. Heatmaps display unique molecular identifier (UMI) counts per spatial location (spot).

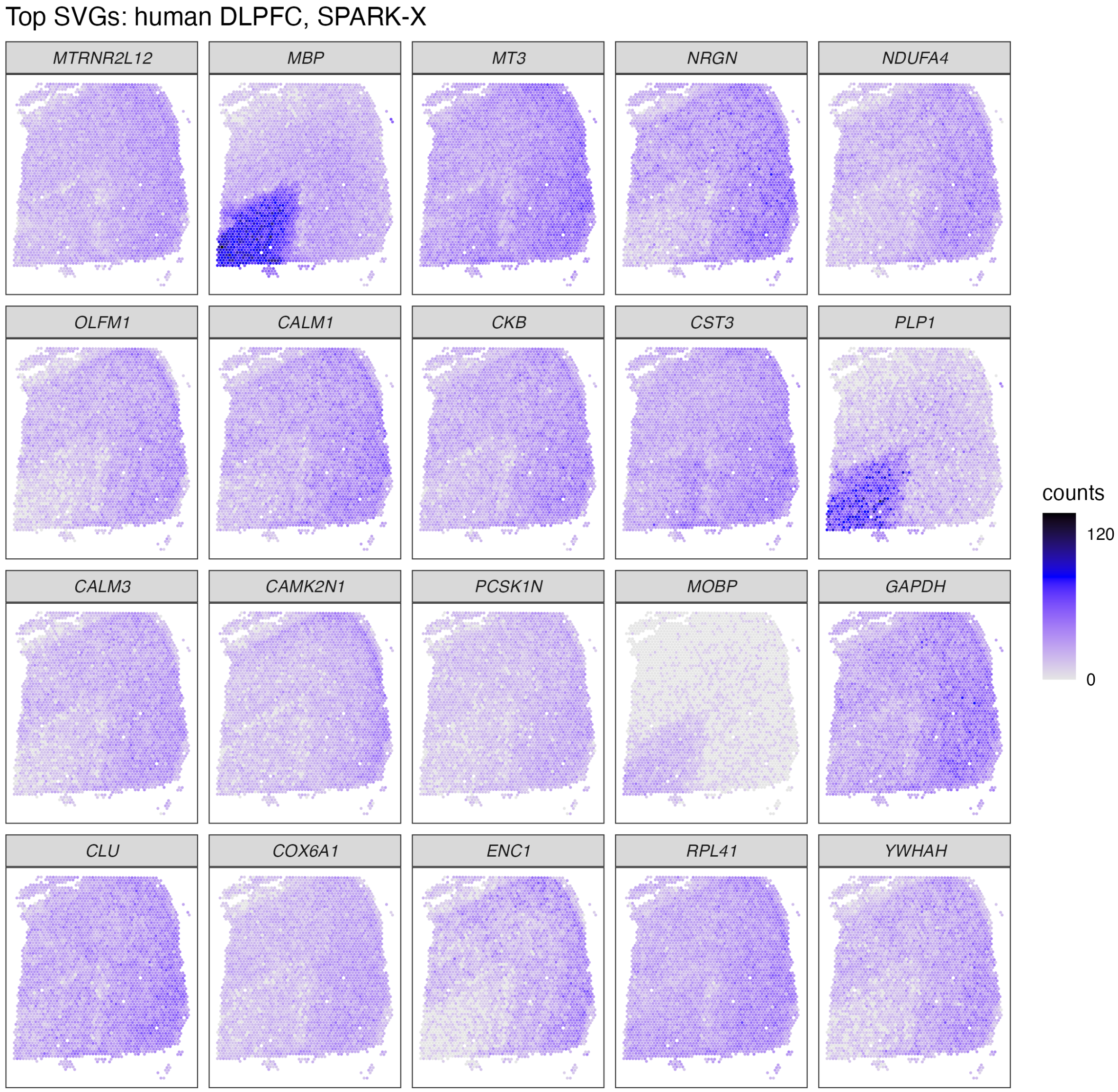


**Supplementary Figure S6: Spatial expression of top SVGs, Visium human DLPFC, SPARK-X.** Spatial expression plots of top 20 SVGs from SPARK-X for Visium human DLPFC dataset. Heatmaps display unique molecular identifier (UMI) counts per spatial location (spot).

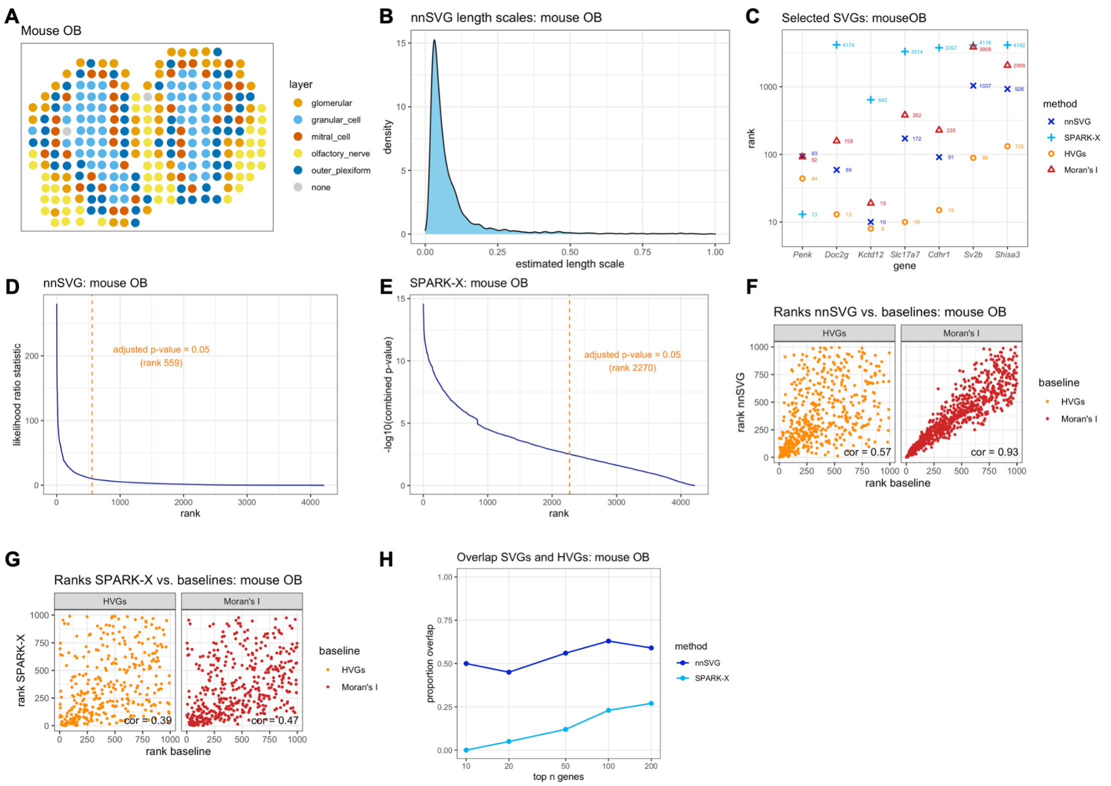

**Supplementary Figure S7: nnSVG recovers biologically informative SVGs in the ST mouse OB dataset.** Using the ST mouse OB dataset [1], nnSVG, SPARK-X, HVGs, and Moran's I were applied to identify SVGs. **(A)** Cell type layer labels per spot with colors representing annotations by [1]. **(B)** Distribution of estimated gene-specific length scale parameters from nnSVG. **(C)** Rank order of 7 known biologically informative SVGs within the lists of top SVGs. **(D)** Estimated likelihood ratio (LR) statistic from nnSVG ($y$-axis) compared to the rank per gene ($x$-axis). Orange dashed vertical line indicates rank cutoff for statistically significant SVGs at a multiple-testing-adjusted $p$-value of 0.05 using LR test with 2 degrees of freedom. **(E)** Combined $p$-value from SPARK-X ($y$-axis) compared to the rank per gene ($x$-axis). Orange dashed vertical line indicates rank cutoff for statistically significant SVGs at a multiple-testing-adjusted p-value of 0.05 using statistical test defined in [27]. **(F)** Ranks of top 1000 SVGs from nnSVG ($y$-axis) compared to ranks from baseline methods ($x$-axis) using HVGs (nonspatial baseline method, left) and Moran's I (spatially-aware baseline method, right), with Spearman correlation (text labels). **(G)** Same as (F) for SPARK-X. **(H)** Overlap between sets of top $n$ SVGs and HVGs ($n = 10, 20, 50, 100, 200$) for nnSVG and SPARK-X.

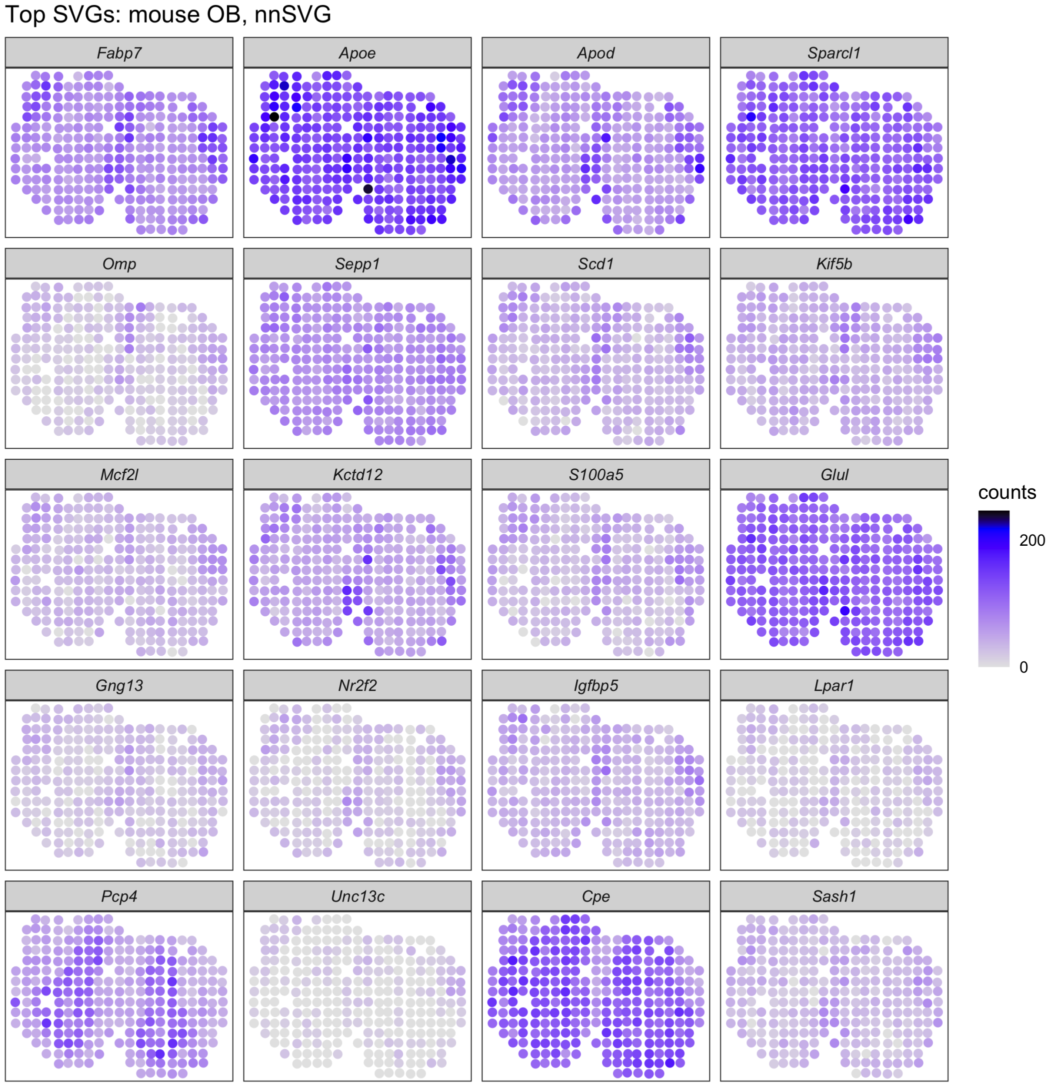


**Supplementary Figure S8: Spatial expression of top SVGs, ST mouse OB, nnSVG.** Spatial expression plots of top 20 SVGs from nnSVG for ST mouse OB dataset. Heatmaps display unique molecular identifier (UMI) counts per spatial location (spot).

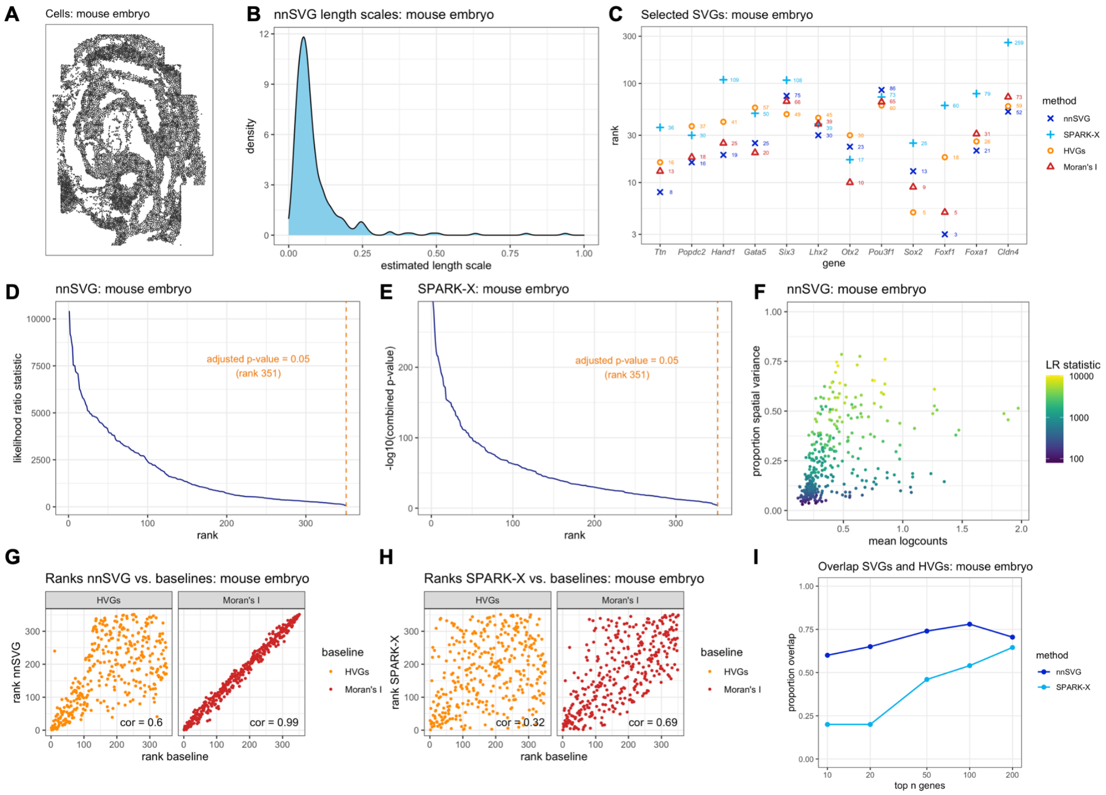

**Supplementary Figure S9: nnSVG recovers biologically informative SVGs in the seqFISH mouse embryo dataset.** Using the seqFISH mouse embryo dataset [38], nnSVG, SPARK-X, HVGs, and Moran's I were applied to identify SVGs. **(A)** Spatial coordinates of cells in this dataset [38]. **(B)** Distribution of estimated gene-specific length scale parameters from nnSVG. **(C)** Rank order of 12 known biologically informative SVGs within the lists of top SVGs. **(D)** Estimated likelihood ratio (LR) statistic from nnSVG ($y$-axis) compared to the rank per gene ($x$-axis). Orange dashed vertical line indicates rank cutoff for statistically significant SVGs at a multiple-testing-adjusted $p$-value of 0.05 using LR test with 2 degrees of freedom. **(E)** Combined $p$-value from SPARK-X ($y$-axis) compared to the rank per gene ($x$-axis). Orange dashed vertical line indicates rank cutoff for statistically significant SVGs at a multiple-testing-adjusted p-value of 0.05 using statistical test defined in [27]. **(F)** Estimated effect size (proportion of spatial variance) along $y$-axis compared to the mean log-transformed normalized counts (logcounts) along $x$-axis for top 1000 SVGs from nnSVG, with the estimated LR statistic per gene indicated with color scale. **(G)** Ranks of top 1000 SVGs from nnSVG ($y$-axis) compared to ranks from baseline methods ($x$-axis) using HVGs (nonspatial baseline method, left) and Moran's I (spatially-aware baseline method, right), with Spearman correlation (text labels). **(H)** Same as (G) for SPARK-X. **(I)** Overlap between sets of top $n$ SVGs and HVGs ($n = 10, 20, 50, 100, 200$) for nnSVG and SPARK-X.

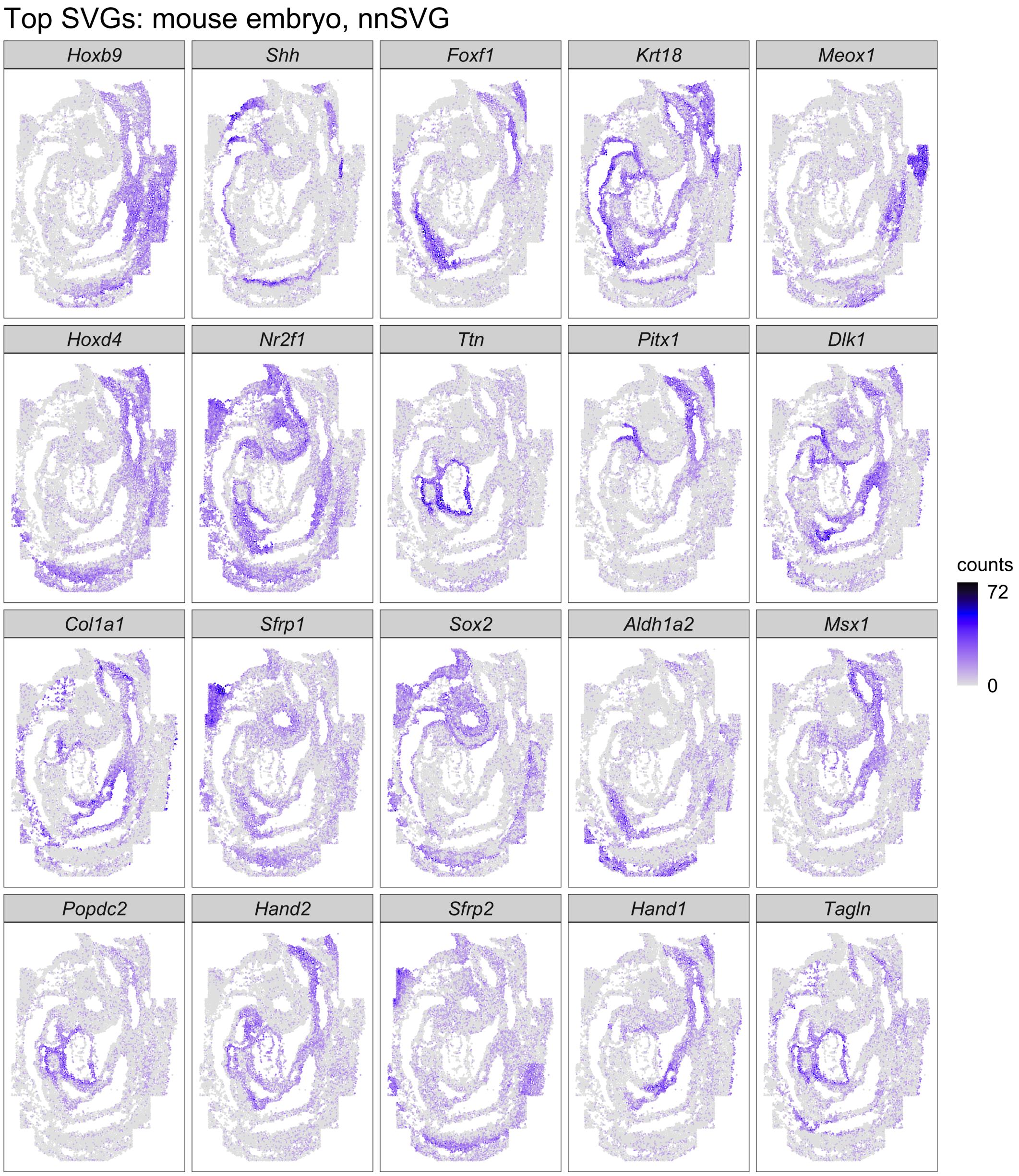


**Supplementary Figure S10: Spatial expression of top SVGs, seqFISH mouse embryo, nnSVG.** Spatial expression plots of top 20 SVGs from nnSVG for seqFISH mouse embryo dataset. Heatmaps display mRNA counts per cell.

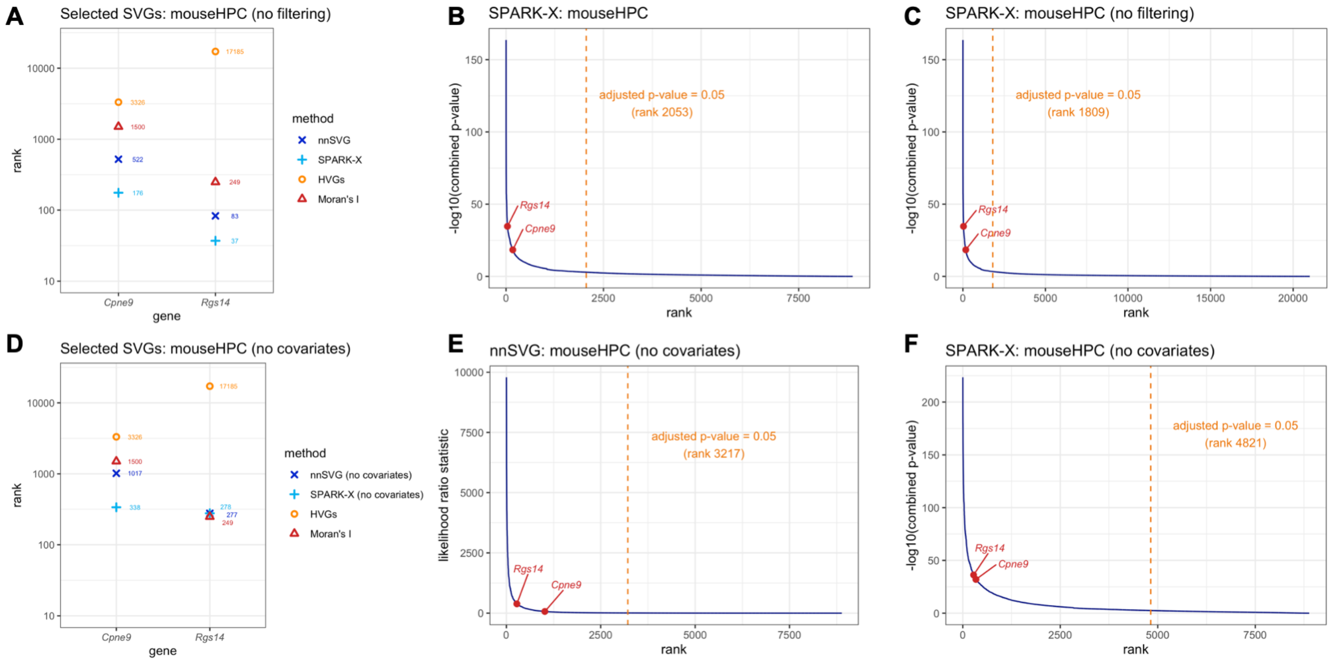


**Supplementary Figure S11: Additional results for Slide-seqV2 mouse HPC dataset. (A)** Rank order of the 2 SVGs from Figure 2 within the lists of top SVGs for nnSVG, SPARK-X, HVGs, and Moran's I, without filtering low-expressed genes for nnSVG or SPARK-X. **(B)** Combined $p$-value from SPARK-X ($y$-axis) compared to the rank per gene ($x$-axis), with the 2 SVGs from (A) labeled. Orange dashed vertical line indicates rank cutoff for statistically significant SVGs at a multiple-testing-adjusted p-value of 0.05 using statistical test defined in [27]. **(C)** Same as (B), without filtering low-expressed genes. **(D)** Rank order of the 2 SVGs from Figure 2 within the lists of top SVGs for nnSVG, SPARK-X, HVGs, and Moran's I, using alternative models for nnSVG and SPARK-X that do not include covariates for spatial domains. **(E)** Estimated likelihood ratio (LR) statistic from nnSVG ($y$-axis) compared to the rank per gene ($x$-axis), using the models for nnSVG that do not include covariates for spatial domains. Orange dashed vertical line indicates rank cutoff for statistically significant SVGs at a multiple-testing-adjusted $p$-value of 0.05 using LR test with 2 degrees of freedom. **(F)** Combined $p$-value from SPARK-X ($y$-axis) compared to the rank per gene ($x$-axis), using the models for SPARK-X that do not include covariates for spatial domains. Orange dashed vertical line as in (B). For the results without filtering low-expressed genes, we used parameters `order = Sum_coords` and `n.neighbors = 15` for nnSVG for improved stability for low-expressed genes.

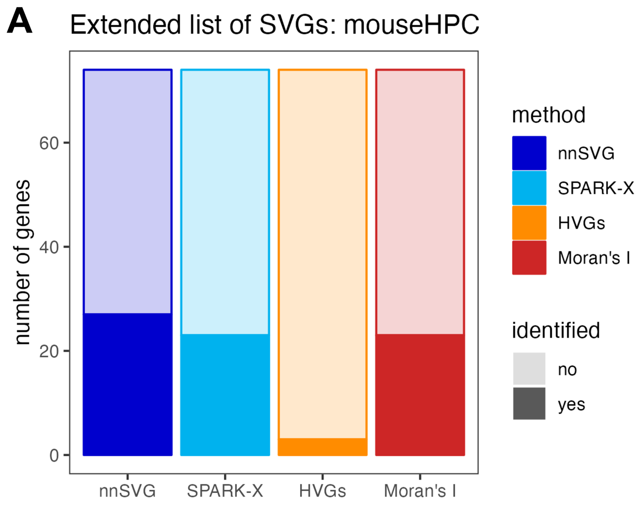


**Supplementary Figure S12: Additional results for Slide-seqV2 mouse HPC dataset. (A)** Number of genes identified within top 1000 SVGs or HVGs for each method, for extended list of 74 known SVGs within the spatial domain defined by CA3 cell type labels, from prior analyses of this dataset [39].

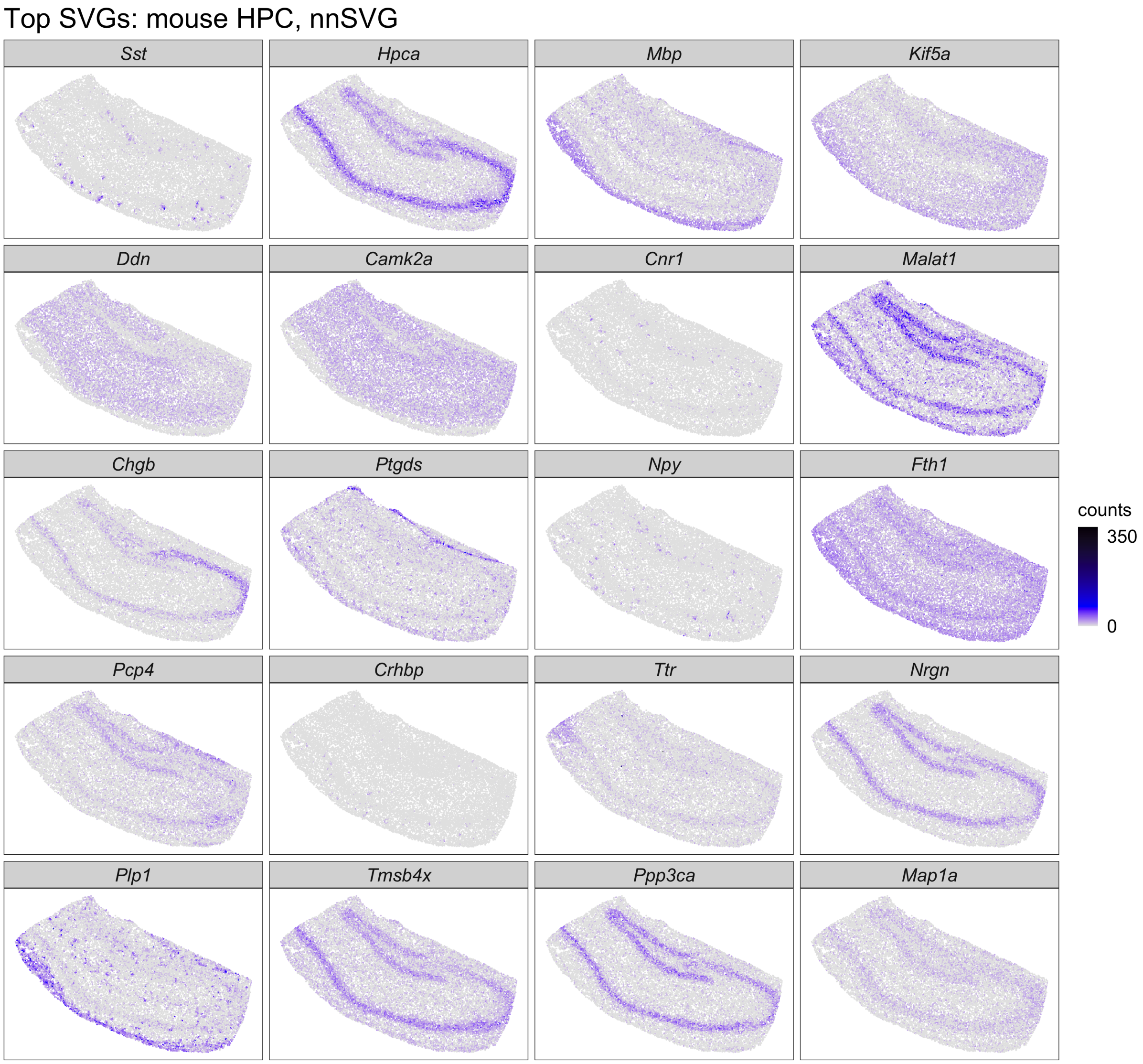


**Supplementary Figure S13: Spatial expression of top SVGs, Slide-seqV2 mouse HPC, nnSVG.** Spatial expression plots of top 20 SVGs from nnSVG for Slide-seqV2 mouse HPC dataset. Heatmaps display unique molecular identifier (UMI) counts per spatial location (spot).

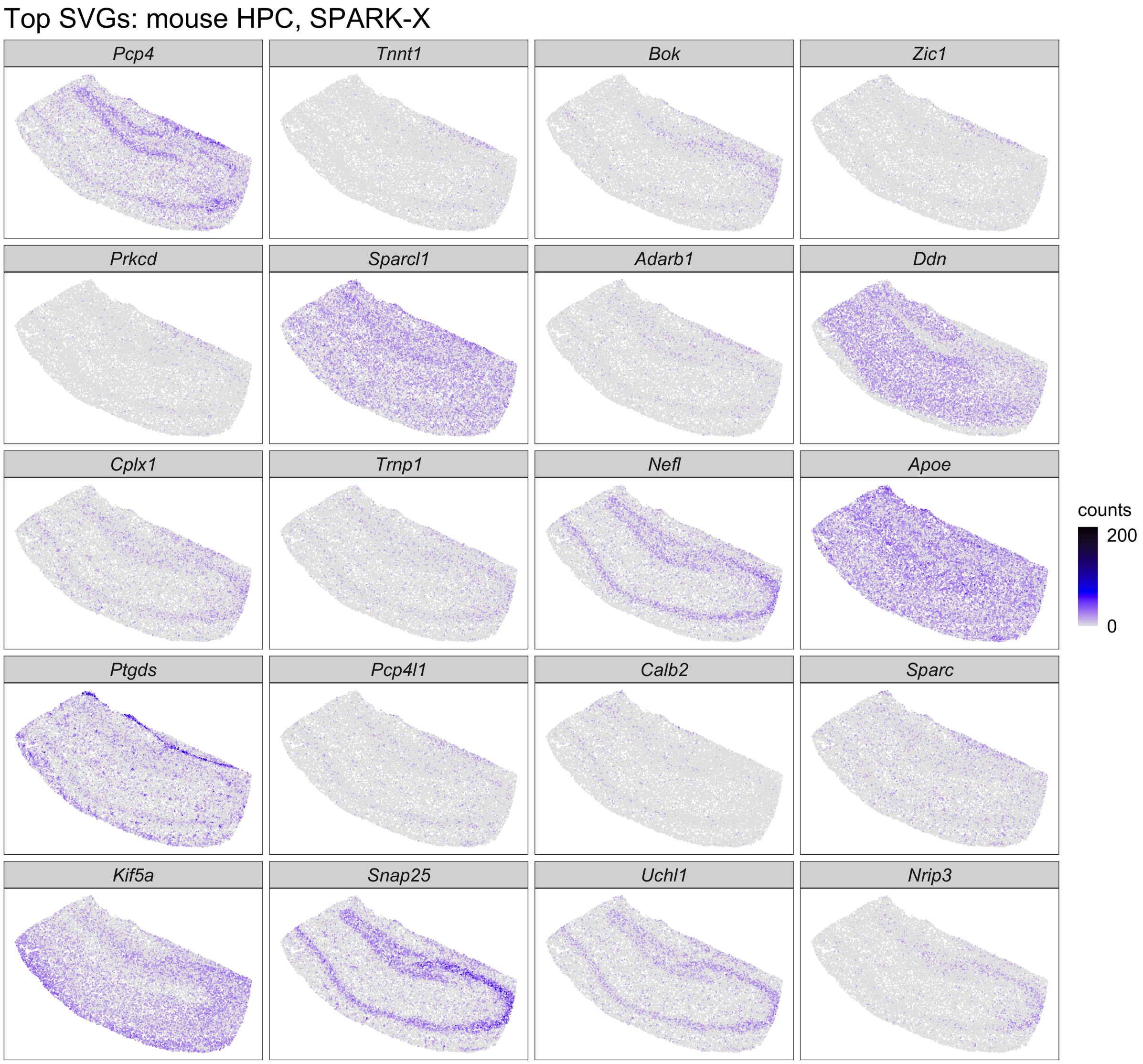


**Supplementary Figure S14: Spatial expression of top SVGs, Slide-seqV2 mouse HPC, SPARK-X.** Spatial expression plots of top 20 SVGs from SPARK-X for Slide-seqV2 mouse HPC dataset. Heatmaps display unique molecular identifier (UMI) counts per spatial location (spot).

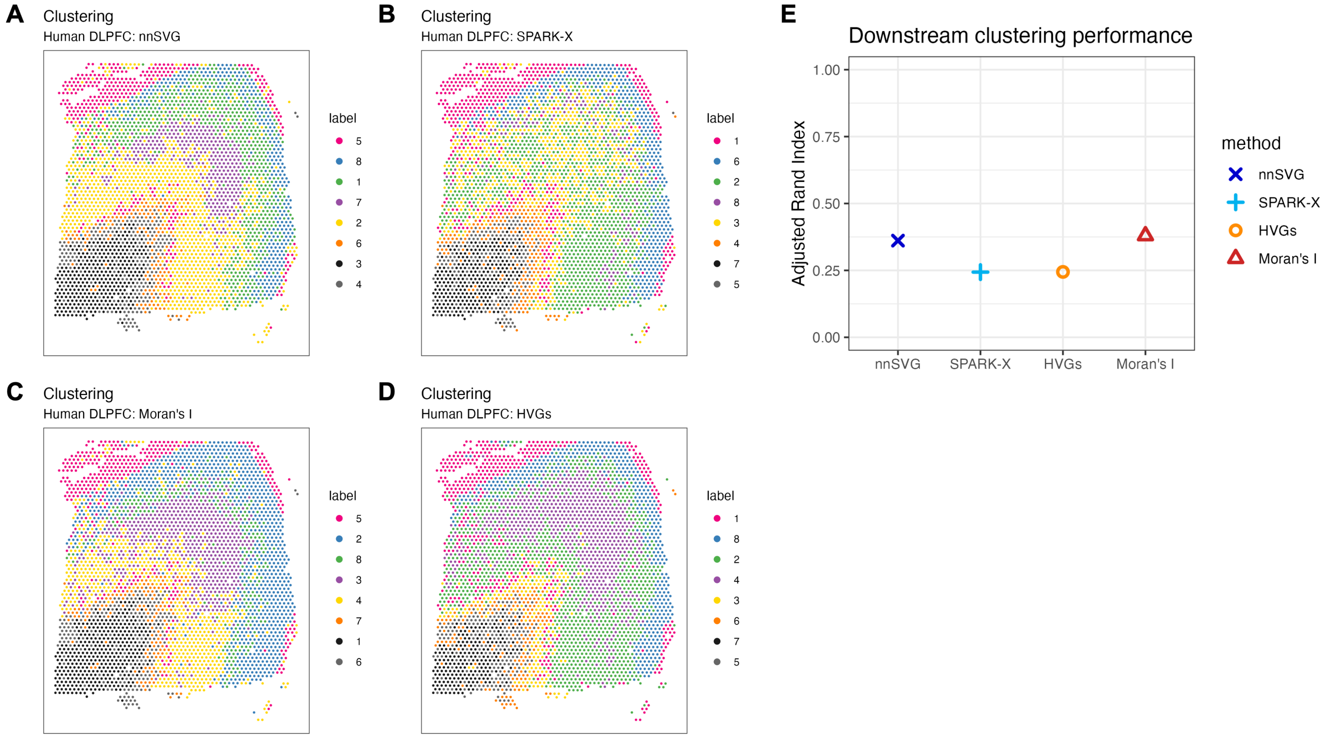


**Supplementary Figure S15: Downstream clustering performance, Visium human DLPFC dataset.** (**A**–**D**) Cluster labels for nnSVG, SPARK-X, Moran's I, and HVGs when using the top 1000 SVGs (nnSVG, SPARK-X, and Moran's I) or top 1000 HVGs as the input for graph-based unsupervised clustering. **(E)** Adjusted Rand index evaluating the similarity between cluster labels and the manually annotated layer labels [8] in this dataset.

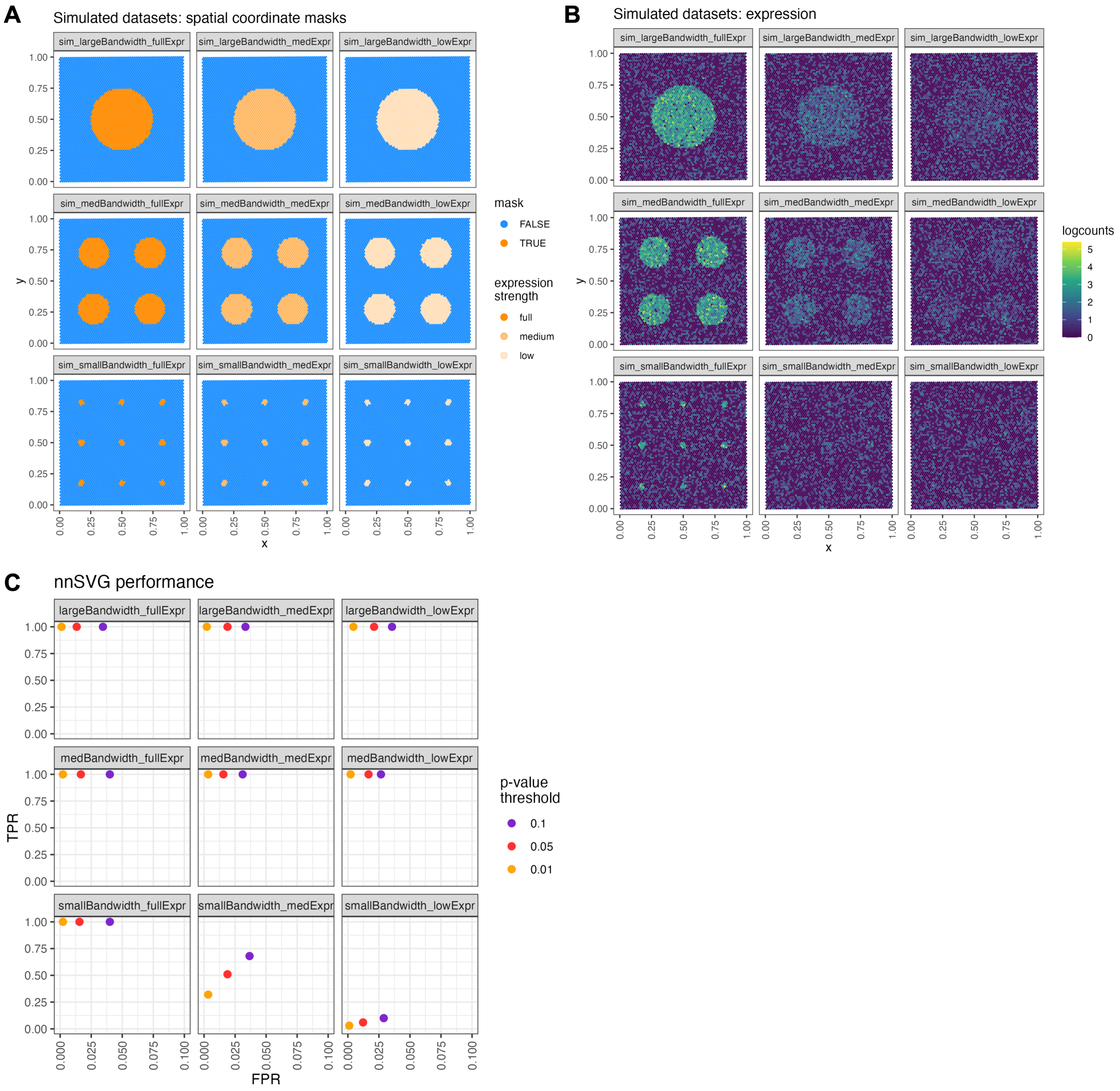


**Supplementary Figure S16: Summary of nnSVG simulations and results.** **(A)** Spatial coordinate masks and **(B)** expression values (log-transformed normalized counts, logcounts) for the main simulations, which consist of scenarios with varying length scales (panels top to bottom) and expression strength (panels left to right) for the simulated SVGs. Each simulation scenario included 100 simulated true SVGs and 900 noise genes. **(C)** Performance evaluations in terms of the true positive rate (TPR) and false positive rate (FPR) for recovering the set of SVGs within each simulation scenario, at nominal $p$-value thresholds of 0.01, 0.05, and 0.1. TPR is defined as the proportion of simulated true SVGs that are correctly identified as statistically significant SVGs, and FPR is defined as the proportion of noise genes that are incorrectly identified as statistically significant SVGs, using likelihood ratio (LR) test with 2 degrees of freedom, at the nominal $p$-value thresholds indicated.

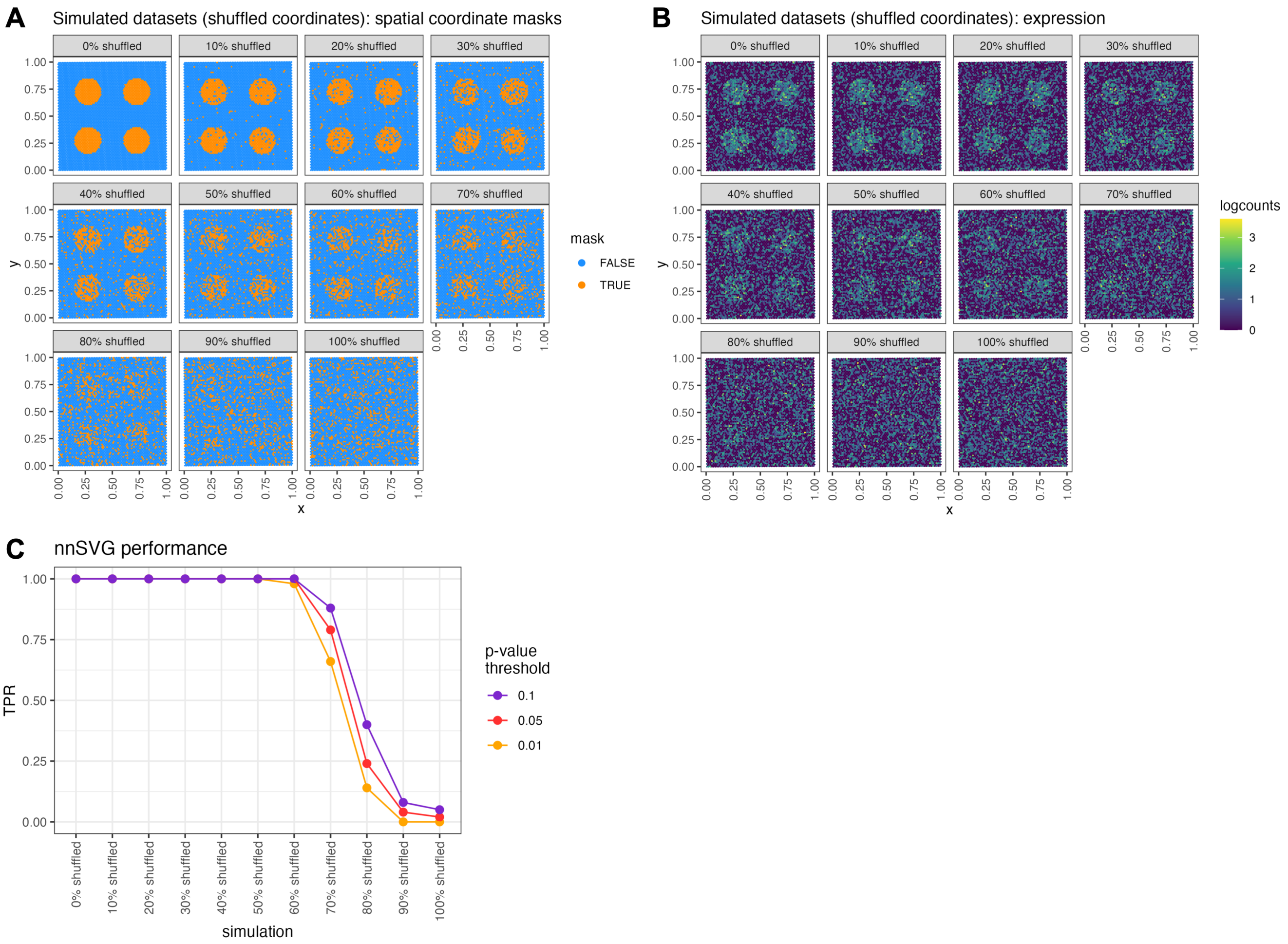


**Supplementary Figure S17: Summary of nnSVG simulations and results.** **(A)** Spatial coordinate masks and **(B)** expression values (log-transformed normalized counts, logcounts) for the shuffled simulations, which consist of scenarios with increasing proportions of spatial coordinates (10%, 20%, ..., 100%) randomly shuffled to introduce noise into the simulations. The shuffled scenarios are based on the 'medium length scale, medium expression strength' scenario from the main simulations in Supplementary Figure S16. **(C)** Performance evaluations in terms of the true positive rate (TPR) for recovering the set of SVGs within each simulation scenario, at nominal *p*-value thresholds of 0.01, 0.05, and 0.1. TPR is defined as the proportion of simulated true SVGs that are correctly identified as statistically significant SVGs, using likelihood ratio (LR) test with 2 degrees of freedom, at the nominal *p*-value thresholds indicated.

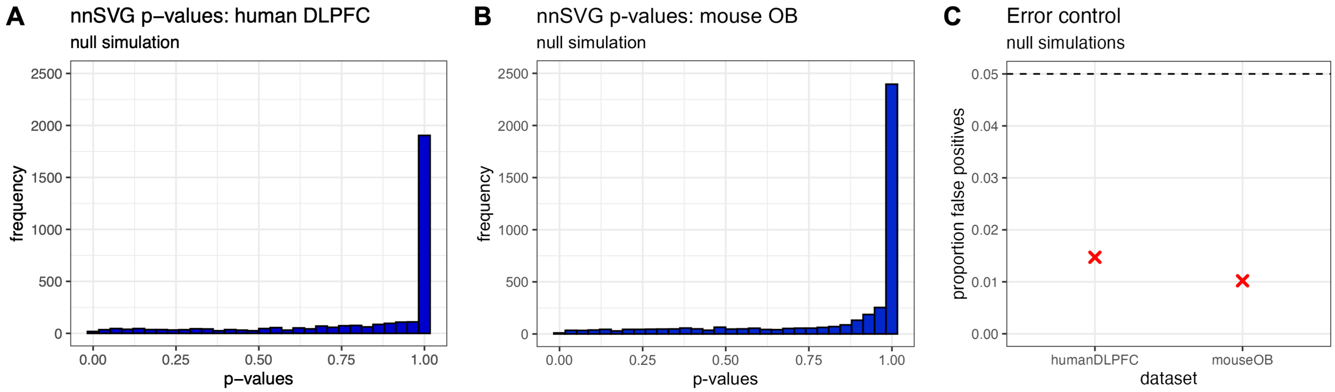


**Supplementary Figure S18: nnSVG null simulations.** **(A–B)** $P$-value distributions from nnSVG for null simulations, where the spatial coordinates were permuted to remove any spatial correlation structure, for **(A)** the Visium human DLPFC and **(B)** the ST mouse OB datasets. **(C)** Proportion of observed false positives at a $p$-value cutoff of 0.05 ($p \leq 0.05$) among the null simulation $p$-values from (A) and (B). Figures show unadjusted $p$-values from likelihood ratio (LR) test with 2 degrees of freedom.

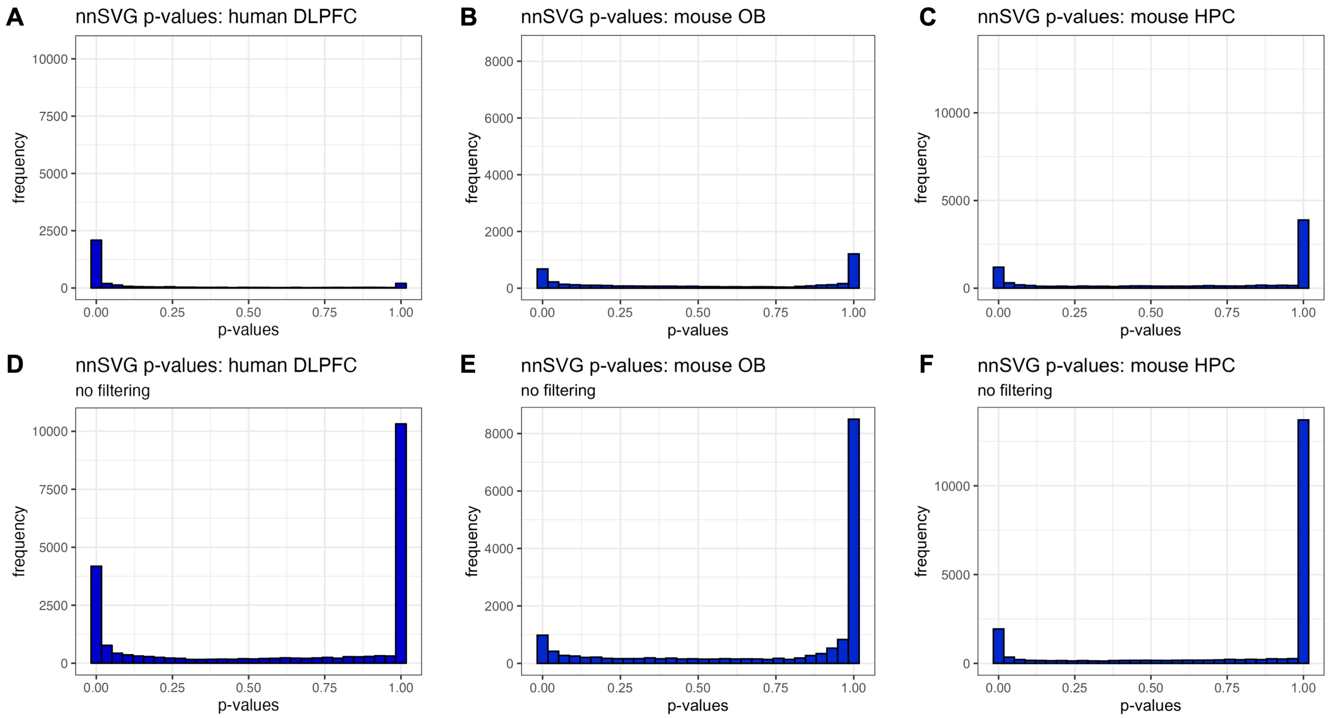


**Supplementary Figure S19: nnSVG *p*-value distributions.** $P$-value distributions from nnSVG for the main results (including filtering of low-expressed genes) for the transcriptome-wide datasets (Visium human DLPFC, ST mouse OB, and Slide-seqV2 mouse HPC) **(A–C)**, and for alternative results without filtering low-expressed genes **(D–F)**. Figures show unadjusted $p$-values from likelihood ratio (LR) test with 2 degrees of freedom.

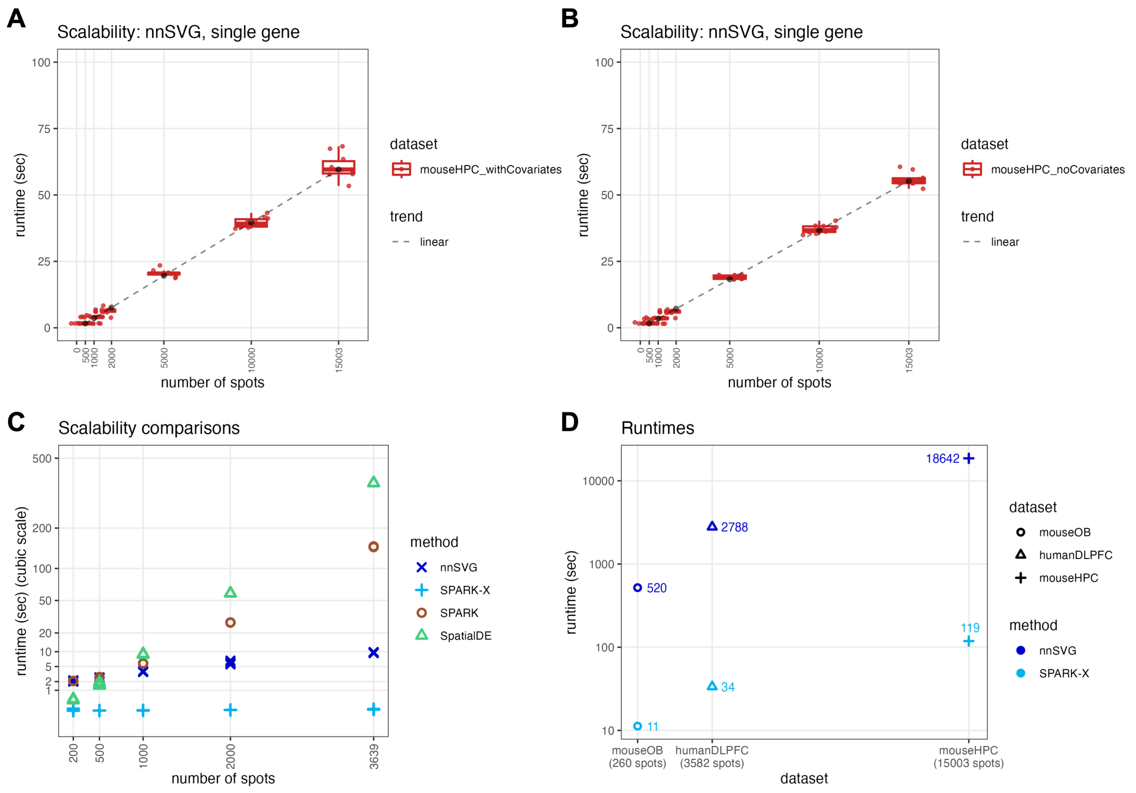


**Supplementary Figure S20: Additional scalability and runtime comparisons. (A–B)** We evaluated the scalability of nnSVG for subsampled numbers of spatial locations in the Slide-seqV2 mouse HPC dataset with covariates **(A)** and without covariates **(B)** included for spatial domains within the models, for the set of annotated spatial locations in this dataset. **(C)** Scalability comparisons across methods, including cubically scaling methods (SpatialDE and SPARK). We subsampled from the full number of spatial locations in the Visium human DLPFC dataset ($n$ = 200, 500, 1000, 2000, and all 3639 spatial locations) and ran each method $n = 10$ times with different random seeds, for two genes (note that SPARK and SPARK-X require at least two genes to run without error), for each number of spatial locations. Note $y$-axis is on cubic scale. **(D)** Runtime in seconds ($y$-axis) for nnSVG and SPARK-X for the three transcriptome-wide datasets ($x$-axis) from the main results, with text labels and axis scale along $x$-axis indicating the number of spatial locations within each dataset (after quality control and filtering unannotated spots). We used 10 processor cores for all datasets for nnSVG and the Slide-seqV2 mouse HPC dataset for SPARK-X, and a single processor core for the Visium human DLPFC and ST mouse OB datasets for SPARK-X. Boxplots show medians, first and third quartiles, and whiskers extending to the furthest values no more than 1.5 times the interquartile range from each quartile.

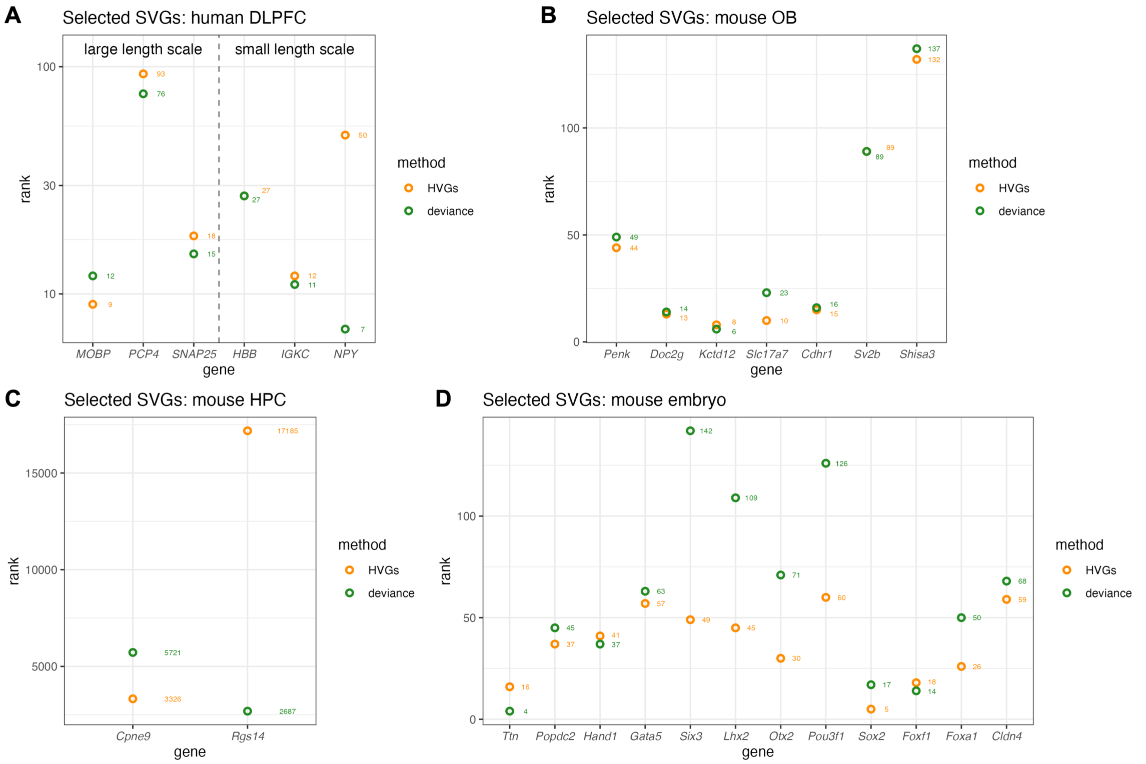


**Supplementary Figure S21: Deviance residuals from binomial model as baseline method.** As an alternative nonspatial baseline method, we compared deviance residuals from a binomial model [20] against the HVGs baseline method for the **(A)** Visium human DLPFC, **(B)** ST mouse OB, **(C)** Slide-seqV2 mouse HPC, and **(D)** seqFISH mouse embryo datasets. For each dataset, we compared the two baseline methods in terms of the rank order of the selected SVGs evaluated in the main results. In the main results, we show HVGs as the nonspatial baseline method.

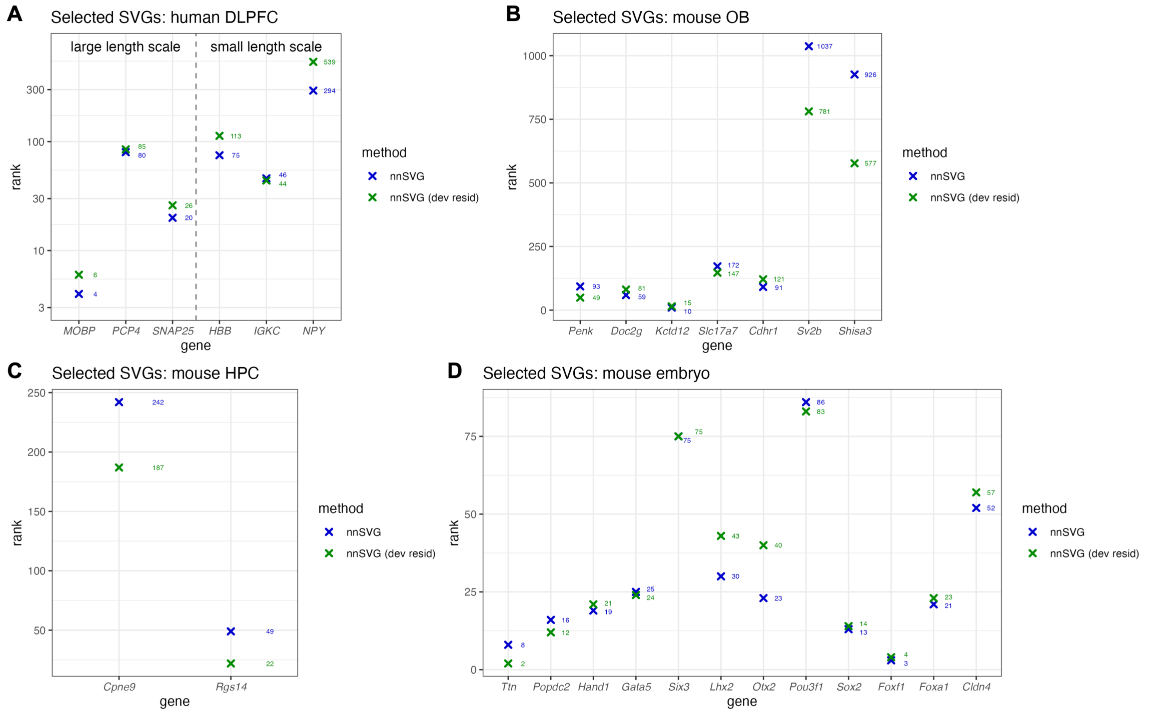


**Supplementary Figure S22: Deviance residuals from binomial model for preprocessing for nnSVG.** As an alternative preprocessing method for the raw data, we compared the results from nnSVG using deviance residuals from a binomial model [20] for preprocessing against the main results using log-transformed normalized counts (logcounts) for preprocessing. We used the `scry` R/Bioconductor package [20] to calculate the deviance residuals, and compared the results from nnSVG using the two preprocessing methods for the **(A)** Visium human DLPFC, **(B)** ST mouse OB, **(C)** Slide-seqV2 mouse HPC, and **(D)** seqFISH mouse embryo datasets. In the main results, we show the results for nnSVG using the logcounts preprocessing method.

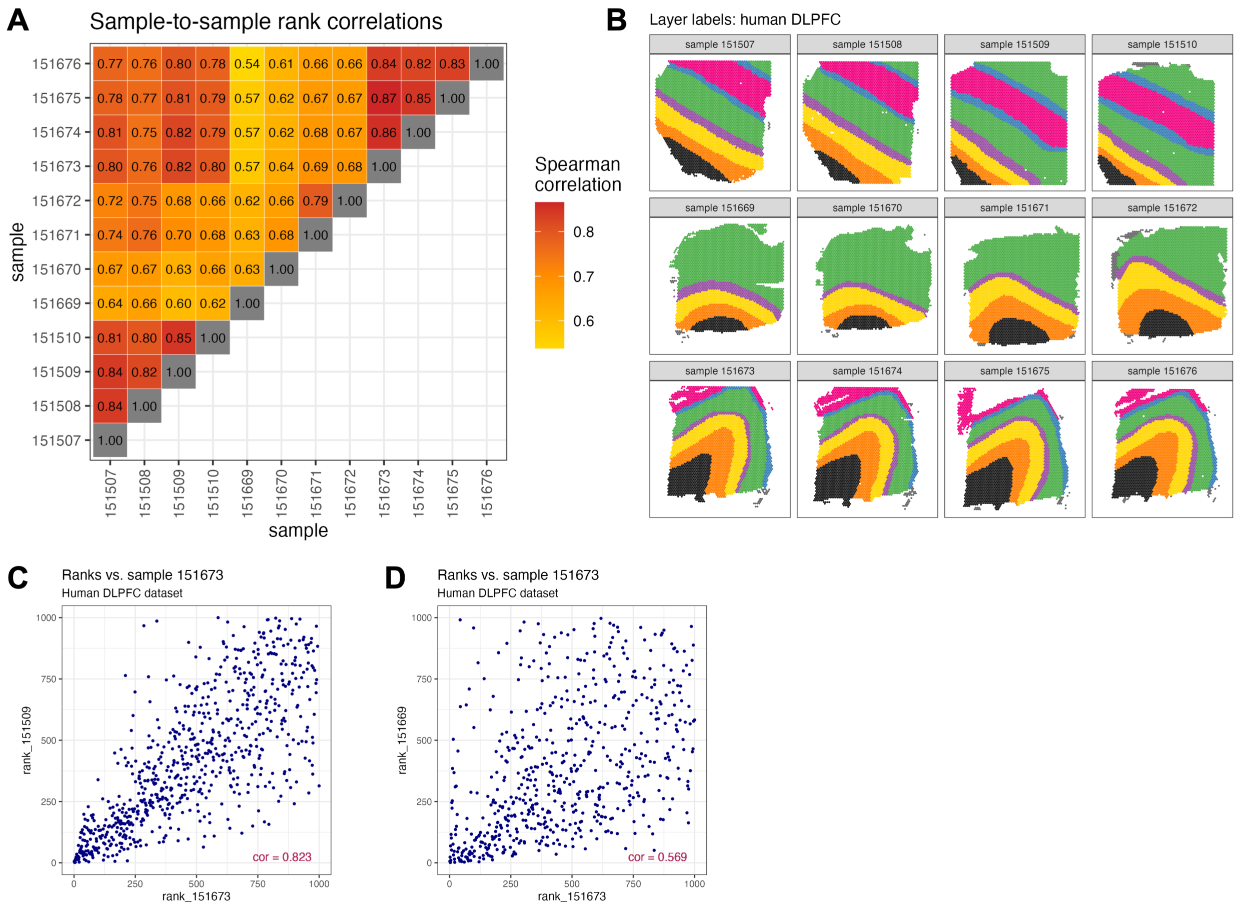


**Supplementary Figure S23: Comparison of nnSVG results across multiple samples.** **(A)** Pairwise Spearman correlations between rankings of SVGs from nnSVG between each pair of samples in the original source of the Visium human DLPFC dataset (12 samples from 3 donors) [8, 43]. Samples from each donor are arranged in blocks of rows and columns (4 samples per donor). **(B)** Manually annotated cortical layer labels for the 12 samples in this dataset, from original source [8, 43]. Samples from each donor are arranged in rows (donors 1 to 3, top to bottom, 4 samples per donor). **(C–D)** Rank comparisons for the samples with the highest **(C)** and lowest **(D)** correlations with sample 151673 (the sample used in the main results). Spearman correlations are shown in text labels.

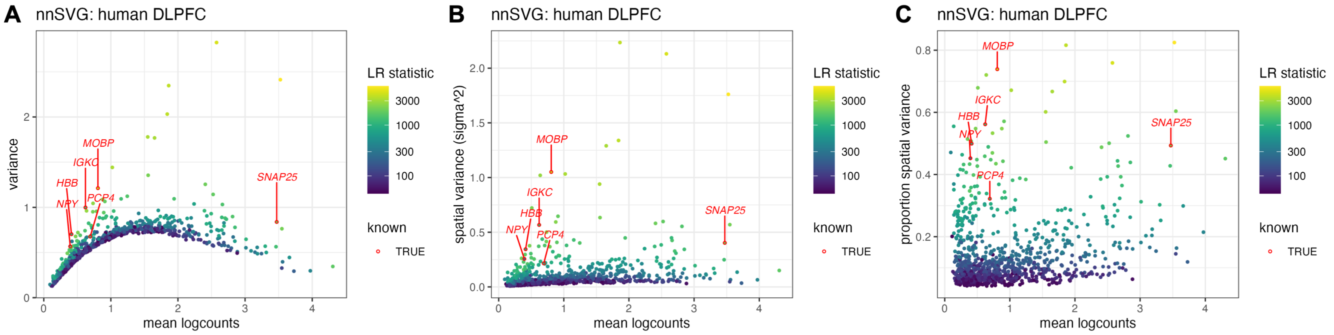


**Supplementary Figure S24: Mean-variance relationship and estimated effect sizes for Visium human DLPFC dataset.** Alternative definitions of effect size vs. mean log-transformed normalized counts (logcounts). **(A)** Total variance vs. mean logcounts per gene, analogous to variance vs. mean logcounts in scRNA-seq workflows, with 6 SVGs from Figure 1 labeled, and estimated LR statistics from nnSVG indicated with color scale. In scRNA-seq workflows, HVGs are ranked by excess 'biological variance' above the mean-variance trend, where the trend is assumed to represent technical variance [17]. **(B)** Spatial variance (estimated $\sigma^2$ from nnSVG) vs. mean logcounts per gene. **(C)** Proportion spatial variance (out of total variance, from nnSVG, see Methods) vs. mean logcounts per gene.
